# Characterization of interindividual DNA methylation variability in rainbow trout (*Oncorhynchus mykiss*)

**DOI:** 10.1101/2025.08.13.669306

**Authors:** Lefort Gaëlle, Brionne Aurélien, Piégu Benoît, Terrier Frédéric, Pigeon Antoine, Beauclair Linda, Roy Jérôme, Marandel Lucie, Panserat Stéphane, Lallias Delphine, Coustham Vincent

**Affiliations:** INRAE, CNRS, Université de Tours, PRC, 37380, Nouzilly, France; INRAE, UR1037 LPGP, Fish Physiology and Genomics, Campus de Beaulieu, F-35000 Rennes, France; INRAE, Université de Pau & Pays de L’Adour, NuMeA, E2S UPPA, Aquapôle, 64310, Saint-Pée-Sur-Nivelle, France; Université Paris-Saclay, UMR 9019 CNRS Intégrité du génome et cancer, Institut Gustave Roussy; INRAE, Université Paris-Saclay, AgroParisTech, Institute Jean-Pierre Bourgin for Plant Sciences (IJPB), 78000, Versailles, France; INRAE, Université Paris-Saclay, AgroParisTech, GABI, 78350, Jouy-en-Josas, France

**Keywords:** DNA methylation, interindividual variability, plasticity, rainbow trout

## Abstract

Interindividual epigenetic variability, particularly in DNA methylation, is now recognized as a significant contributor to phenotypic diversity in humans and mammals. These epivariable regions, which make up a small fraction of the genome, are strongly influenced by genetic factors and environmental factors, especially during early development. In this context, epigenetic variability of DNA methylation has been proposed as an adaptive force involved in various environmental responses. In fish and other vertebrates, environmental factors are known to influence the health, performance and welfare, likely through the alteration of the epigenetic landscape. However, whether interindividual epigenetic variability may contribute to the phenotypic plasticity of fishes is unknown. Here we provide a first description of the rainbow trout methylome variability using a whole-genome bisulfite sequencing approach in an isogenic line to minimize genetic variation. Variable methylation regions were identified in both liver and hypothalamus tissues of 12 replicate fishes and were found enriched at gene regulatory elements, such as promoters and first introns. Gene Ontology analysis revealed functional clusters related to cellular development, neural communication, metabolic balance, and immune response. Interestingly, some variably methylated regions are found at the same genomic loci in both tissues and showed a strong intraindividual correlation in methylation levels, suggesting establishment during early embryogenesis. Overall, our work demonstrates the existence of interindividual epigenetic variability in rainbow trout and provides valuable insights into the regulatory function of DNA methylation variation that is likely involved in developmental and physiological processes.

## Introduction

Epigenetics is commonly defined as the set of molecular mechanisms involved in gene expression regulation that are reversible and transmissible during development, and sometimes between generations, without altering the DNA sequence. The epigenome (the whole set of epigenetic marks in a tissue) responds to environmental fluctuations such as temperature or nutrition and can be considered as a mediator between genes and the environment [1]. In this context, environmental programming strategies during a window of phenotypic plasticity (often during early development) have successfully been used as a lever to adjust the phenotype of individuals to their living conditions [2]. However, the mechanisms underlying phenotypic variation are still poorly understood, and the epigenome-wide identification of environmental exposure is often hindered by interindividual variability.

Interestingly, interindividual epigenetic variability may not just be noise, but may also contribute to the animal’s response to stimuli from the environment. Indeed, the work of Feinberg and Irizarry [3] demonstrated the existence of variable methylation regions in mice and humans, referred to as VMRs (Variable Methylation Regions). VMRs represent clusters of CpGs with highly polymorphic DNA methylation levels in a population and were shown to be enriched for various functional genomic features, in particular enhancers, suggesting a role in gene expression regulation [4]. VMR mapping in the human genome has shown that these epigenetic variations are due to both genetic and environmental causes [4]. Further studies showed that VMRs are also found in cattle, sharing the same characteristics [5]. Therefore, VMRs appear to be non-randomly distributed, conserved in several species of mammals and potentially important for perceiving and/or remembering the early effects of the environment.

Based on functional analysis, it was proposed that VMRs could contribute to phenotypic plasticity in changing environments [3]. In this context, understanding the contribution of interindividual epigenetic variability (VMRs) to phenotypic plasticity is particularly relevant in fish that are also subjected to important environmental fluctuations. In an agronomic context, epigenetic variability may enhance aquaculture resilience in changing environments by enabling the selection of individuals with epigenetic profiles linked to stress tolerance, disease resistance, and growth efficiency. For instance, hypomethylation of stress-response genes in certain fish correlates with improved adaptation to temperature fluctuations or hypoxia [6]. Epigenetic markers also provide insights into phenotypic plasticity, helping predict how populations might respond to environmental shifts [7]. To explore the existence of VMRs in a fish species, and to gain insights into the functions of these regions, we carried out a characterization of VRMs in an isogenic (*i*.*e*. genetically identical) line of rainbow trout. Observed differences in methylation are therefore more likely to reflect genuine epigenetic variation rather than underlying genetic differences. Liver and hypothalamic trout methylomes were studied by whole-genome bisulfite sequencing (WGBS) in 12 genetically identical individuals. These two organs were chosen because of their importance in regulating metabolism and adapting to stresses [8]. Our study revealed several hundreds of VMRs in both tissues, a quarter of which were common between the tissues, located mainly in introns and intergenic regions, suggesting functional conservation of these regions in teleost fish.

## Methods

### Animals, tissue collection and DNA extraction

The study was conducted on a heterozygous isogenic line, AB1h, that was produced at the PEIMA INRAE experimental fish facilities (IE PEIMA [9], INRAE, Sizun, France, registration number D29-277-02) by crossing several females from the B57 homozygous isogenic line with a single male from homozygous line AB1. Homozygous isogenic lines used in the present study were originally established and are maintained at INRAE [10]. Eggs were transferred at the Lees-Athas INRAE fish facilities for the first month of life (IE NuMéA [11], INRAE, Lees-Athas, France, registration number A64.104.1,) and then transferred at the Donzacq INRAE fish facilities right before the first meal for the remainder (IE NuMéA [11], INRAE, Donzacq, France, registration number A40-228.1).

The trout were fed *ad libitum* twice a day with a standard commercial feed (NUTRA HP, Skretting, Fontaine-les-Vervins, France). The weight of the fish (total biomass per tank) and the quantity of feed distributed were measured every 3 weeks. After 6 months of feeding, the trout were fasted for 72 hours then sedated, anaesthetized and euthanized with growing concentrations (15, 50 and 150 mg/L, respectively) of tricaine methanesulfonate (PharmaQ, Norway) in a water bucket. The 12 animals with the least variation around the mean weight (37.6 g ± 2.58) were selected from 30 animals taken from two replicated tanks. Liver and hypothalamus tissue samples were frozen in liquid nitrogen immediately after sampling. High molecular weight genomic DNA was extracted using MagAttract HMW DNA Kit (Qiagen, Coutarboeuf, France) following manufacturer’s recommendations. DNA integrity was checked on a 0.8 % agarose gel and DNA concentration was determined using a Nanodrop ND-1000 Spectrophotometer (Marshall Scientific, USA).

### WGBS library preparation, sequencing, and data processing

Post-bisulfite adapter tagging whole-genome bisulfite (PBAT-WGBS) library preparation and DNA sequencing were performed by Novogene (UK). 150 bp paired-end libraries were sequenced on the Illumina Novaseq 6000 platform. Data processing and analysis were performed at the ISLANDe technical platform (UMR PRC, Nouzilly, France). Quality control of the sequencing data was performed with [FastQC](https://www.bioinformatics.babraham.ac.uk/projects/fastqc) (v. 0.11.9). Quality and adapter trimming were performed with [Trim Galore!](https://www.bioinformatics.babraham.ac.uk/projects/trim_galore/) (v. 0.6.6) using the following arguments: stringency = 5, clip_R1 and clip_R2 = 10. Bismark [12] version 0.23.0 was used to perform alignments of bisulfite-treated reads to the rainbow trout reference genome [USDA_OmykA_1.1](https://www.ncbi.nlm.nih.gov/datasets/genome/GCF_013265735.2/) (GCA_013265735.3) using default parameters. Data was manipulated with Samtools (v 1.11).

### Variable methylation regions identification and analysis

#### VMR identification

The identification of VMRs was conducted using R (version 4.4.1) [13]. Various coverage thresholds, ranging from 5 to 12, for 10 to 90 % of the samples, were tested for each CpG to assess their impact on the number of the remaining positions (Supplementary Table S3 and Supplementary Figure S1). A minimum coverage threshold of 10 in more than 50 % of the samples was selected based on the number of regions (defined by at least 10 CpG sites separated by a maximum of 1,000 bp) identifiable using these parameters (Supplementary Figure S1B). The variability of CpG methylation level for each position was evaluated by calculating the Median Absolute Deviation (MAD), a robust measure of variability for quantitative data [14]. Variable Methylation Cytosines (VMCs) are defined as the cytosines with the highest MAD values. To identify VMRs, we tested different parameters including the maximum distance between two VMCs (100 or 1,000 base pairs apart), the minimum number of VMCs per VMR (5 or 10), and the MAD quantile from the 90th (Q90) to the 99th (Q99) percentile. In all analyses presented in this article, VMRs correspond to at least 5 VMCs spaced no more than 100 base pairs apart from each other. Finally, VMRs located 200 bp or less apart were merged to avoid fragmentation. The quantile used for the analyses was either Q99 or Q90 depending on the needs of the analyses, to focus on the most significant VMRs or to explore a larger set of VMRs, respectively.

#### VMR analysis

VMRs were considered common between both tissues if their coordinates shared at least one base. Data manipulation was performed using the R packages GenomicRanges and plyranges. For each individual, the correlation of VMR levels between the two tissues was calculated for all the CpG positions, by comparing the correlation values between the common regions and the non-common (= tissue specific) regions. The correlation analysis of VMRs and histone mark peaks, ATAC peaks reflecting DNA accessibility and SE (Super-Enhancer) regions obtained from Additional Files 3 and 4 from [15] were performed using R.

#### VMR visualization

BigWig files were obtained from bam files using bedGraphToBigWig program from ucsc tools (v377, https://github.com/ucscGenomeBrowser/kent). Methylation tracks (BigWig files) were visualized using Integrative Genomics Viewer (IGV) [16] version 2.18.4, and VMR regions were imported from a bed file. All tracks presented in the study display the same data range using the “Group Autoscale” function in IGV.

#### Genomic features analysis

The annotation of VMRs was performed using NCBI gene features. The GenomeFeatures R Package (https://forge.inrae.fr/aurelien.brionne/GenomeFeatures, version 0.99) was used to annotate VMRs and examine their distribution across various genomic features, including promoters (defined as 2,000 bp upstream of the Transcription Start Site), flanking exons (5’UTR, 3’UTR), exons, introns, and downstream regions (1,000 bp from the Transcription End Site) on both strands. Genomic features statistically enriched in VMRs compared to the reference genome were identified using a Fisher’s exact test (greater alternative) based on their CpG content. This approach accounts for the size of different genomic features, the heterogeneity of CpG density across the genome, and the preservation of other genomic characteristics.

#### GO analysis

Biological interpretations were conducted using the Gene Ontology (GO) public database, focusing on the Biological Process (BP) category, with the ViSEAGO R package [17]. The analysis was performed on the 63535 and 4908 VMRs identified at the 90^th^ percentile (each VMR comprising at least 5 VMCs no more than 100 bp apart) for hypothalamus and liver, respectively. Gene terms associated with *Oncorhynchus mykiss* (USDA_OmykA_1.1) were identified using the NCBI gene orthologs dataset (ftp.ncbi.nih.gov/gene/DATA/gene_orthologs.gz) to expand the EntrezGene annotations. Enrichment tests were performed using Fisher’s exact test and the “elim” algorithm for each comparison [18]. All enriched GO terms (p < 0.01) were grouped into functional clusters using hierarchical clustering based on Wang’s semantic similarity, respecting the GO graph topology and Ward’s criterion [17]. The clusters were grouped using hierarchical clustering based on the BMA distance between sets of GO terms and Ward’s criterion.

### Sanger sequencing

The LOC110491659 VMR region was selected for genomic sequencing due to the clear variation in methylation levels (Fig. 1A). The region was amplified from unconverted gDNA from the same 12 individuals used for sequencing using P1 (Forward) and P2 (Reverse) primers (Supplementary Table S7). Primers P3-6 were used for Sanger Sequencing by Azenta Life Sciences (UK). 500 ng of genomic DNA was bisulfite-converted using the EpiJET Bisulfite Conversion Kit (Thermo Fisher Scientific, USA). Bisulfite-specific P7 (Forward) and P8 (Reverse) primers (Supplementary Table S7) were designed in regions not containing CG dinucleotides using MethPrimer software [19]. PCR was performed using Platinum Taq in a total volume of 20 μl with 1.5 mM MgCl2 following manufacturer’s conditions (Invitrogen, Thermo Fisher Scientific, USA). PCR product specificity and concentration were checked on a 2 % agarose gel and then sequenced (Sanger) by Azenta Life Sciences (UK) using the same primers pair. Methylation levels were calculated by measuring relative peak heights in direct bisulfite-PCR sequencing traces as described previously [20].

**Fig. 1:**
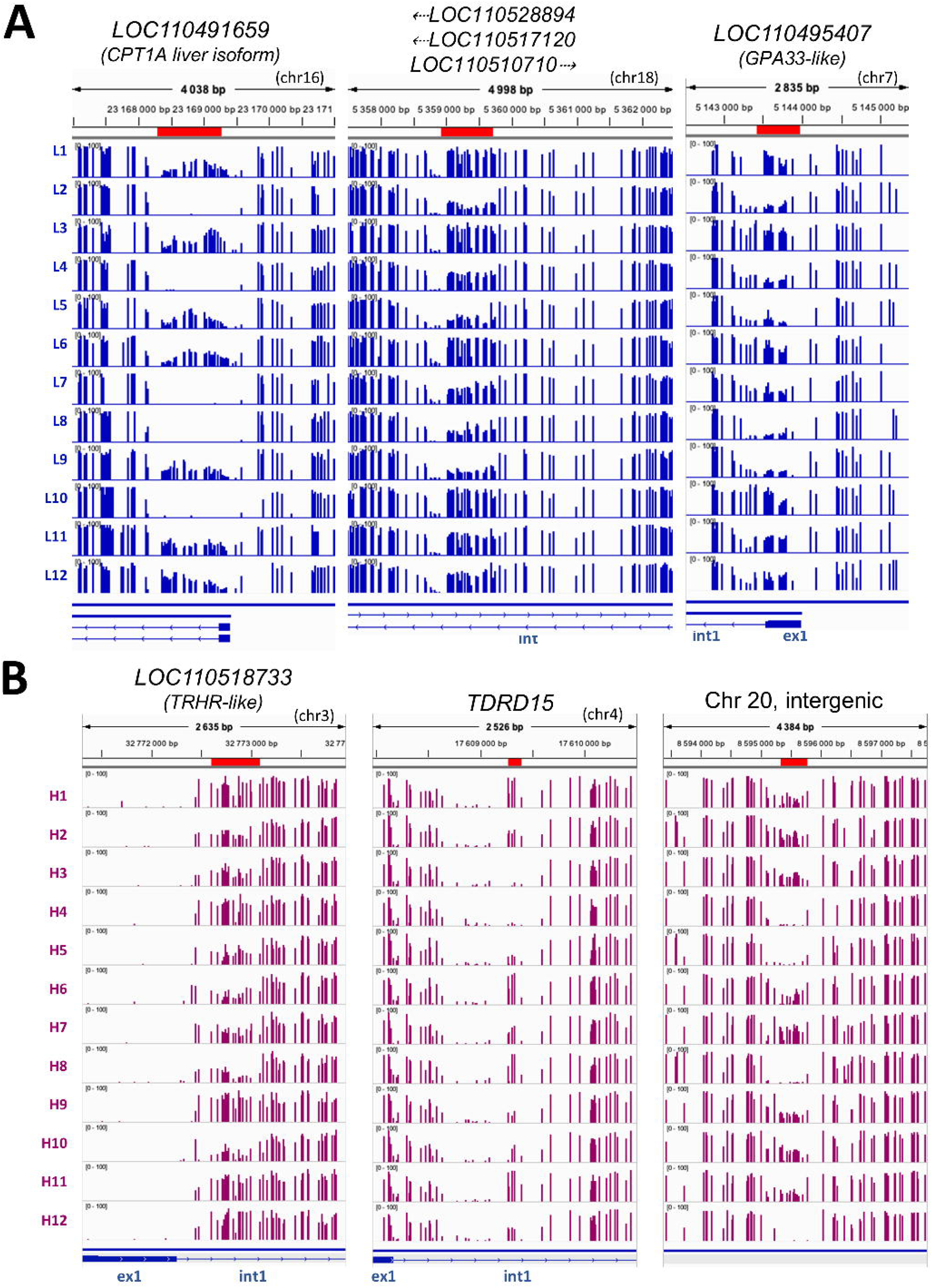
Identification of VMRs. Methylome tracks showing a selection of (A) liver and (B) hypothalamic VMRs for the 12 individuals (blue L1-12 and pink H1-12, respectively). VMR coordinates (Q90) are shown in red boxes. The diagram below the tracks shows the relative positions of exons (plain boxes), introns (narrow arrowed lines), and neighboring genes with directional arrows indicating gene orientation with the exception of VMR in the intergenic region of chromosome 20, where no gene is present.

## Results

### Identification of rainbow trout VMRs

To identify intrinsic DNA methylation variation, rainbow trout were raised in a common, controlled environment. Hepatic and hypothalamic methylomes were obtained from 12 animals. On average, 5 to 6 billion CpGs were analyzed representing a mean coverage of 12.5 and 15.4 X for liver and hypothalamus, respectively (Supplementary Tables S1 and S2) corresponding to ∼7-19 % of total genomic CpGs reaching the minimum coverage threshold (≥ 10) in at least 50 % of samples for both tissues (Supplementary Table S3). VMRs were defined taking into account the maximal distance between VMCs (identified using MAD), the number of VMCs and the percentile of MAD distribution. According to the parameters used and tissues, tens to hundred thousand of VMRs were identified (Supplementary Table S4). Based on these results, we selected two parameters for further analysis that minimize the distance between VMCs while ensuring a sufficient number of VMRs (100 bp and 5 VMCs per VMR, respectively). At the 99th percentile in these conditions, 139 and 814 VMRs were identified for the liver and the hypothalamus, respectively (Supplementary Table S4). A selection of VMRs is illustrated in Fig 1.

To test the reliability of our approach, we used bisulfite conversion followed by Sanger sequencing to analyze in the same individuals the DNA methylation levels at the VMR identified in the first intron of the LOC110491659 gene (Fig. 1A). Although our candidate approach was not as quantitative, we were able to confirm methylation variations among the 12 individuals (Fig. 2). To investigate whether the difference in methylation is directly related to underlying sequence variation or polymorphisms, we performed Sanger sequencing of untreated gDNA of a 1 kb region around the same VMR. As expected by using an isogenic line that is characterized by within-line genetic uniformity, we could not identify *Cis* variation or polymorphisms directly associated with the VMR (Fig. 2).

**Fig. 2:**
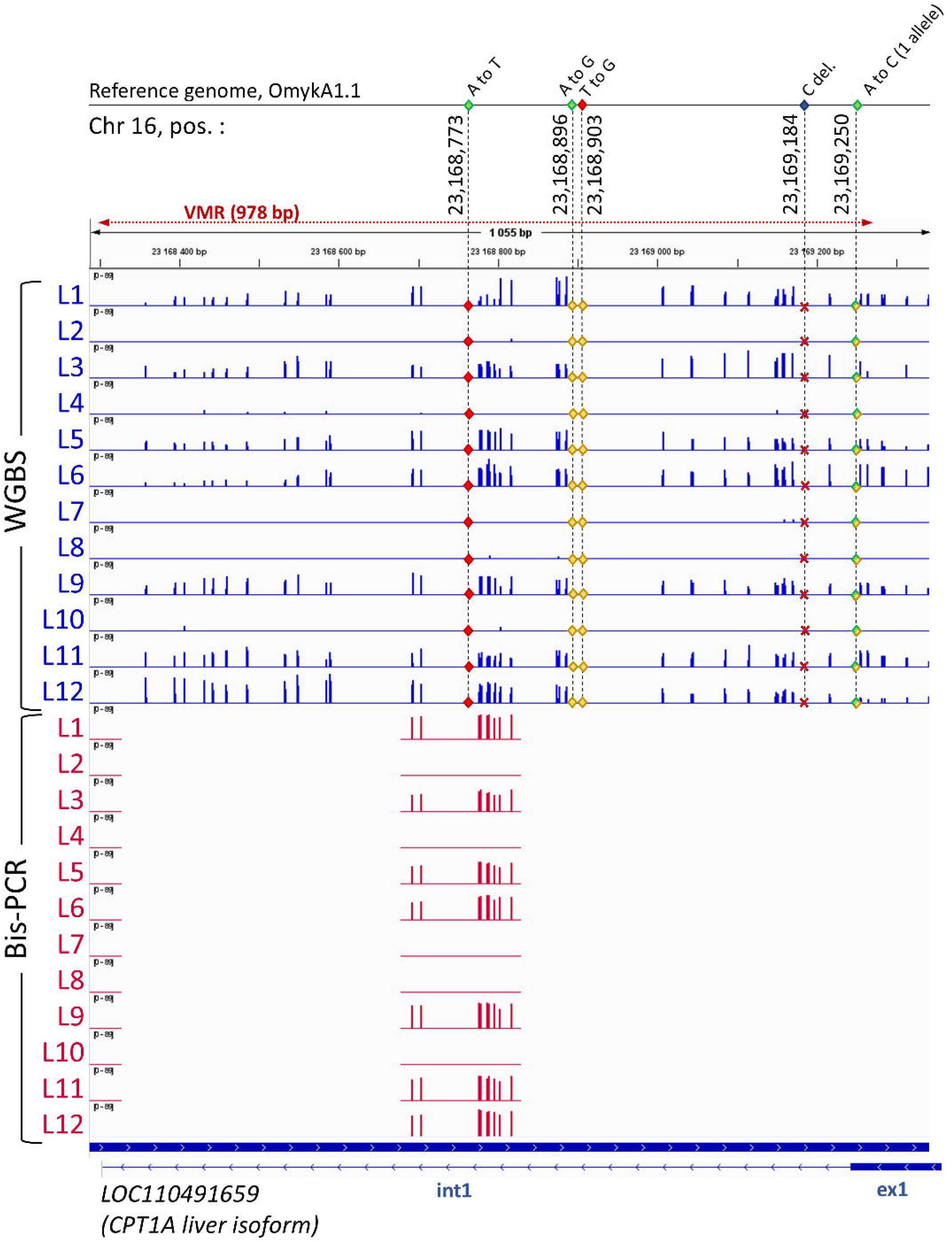
VMR validation in liver and analysis of the underlying genetic variation. The top panel lists specific nucleotide changes (A to T, A to G, T to G, C deletion, A to C in one allele) at positions 23,168,773; 23,168,896; 23,168,903; 23,169,184; and 23,169,250 of chromosome 16 identified in the AB1h isogenic line using Sanger sequencing on genomic DNA compared to the reference genome USDA_OmykA_1.1. No single nucleotide polymorphism was found between the 12 individuals in the 978 bp variable methylated region (VMR) in the CPT1A liver isoform gene (LOC110491659). The red tracks show the % methylation levels identified by bisulfite-PCR followed by Sanger sequencing (Bis-PCR) for the 12 individuals (L1-12) confirming the whole-genome bisulfite sequencing (WGBS) data (blue tracks). The diagram below the tracks shows the relative positions of exons (plain boxes), introns (narrow arrowed lines), and neighboring genes with directional arrows indicating gene orientation.

We then investigated whether some of the VMRs could be common to both tissues. Approximately one quarter of the hepatic VMRs co-localized with the hypothalamic VMRs (Fig. 3A), suggesting genetic or developmental control of these regions.

**Fig. 3:**
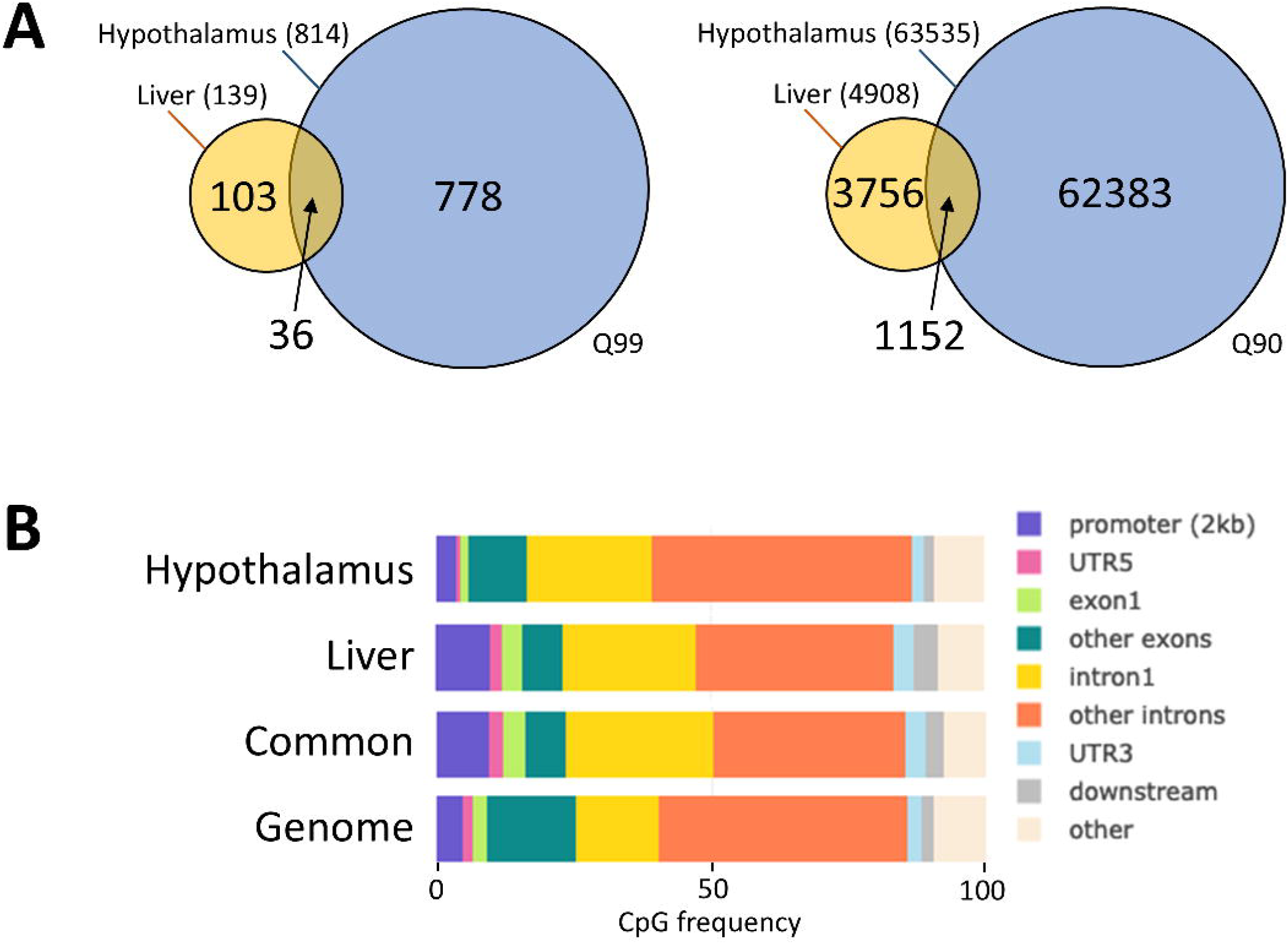
Overlap of VMR between tissues and analysis of genomic distribution. (A) Venn diagram showing the number of VMRs identified for each tissue and the overlap between tissues at Q99 (left) and Q90 (right). (B) Percentage of CpGs contained in VMRs (hypothalamic, hepatic or common between tissues at Q90) or from the whole genome across genomic features, obtained using GenomeFeatures. Promoter and downstream regions were defined upstream of Transcription Start and End Sites (TSS and TES, respectively), respectively.

### VMR distribution suggests a gene regulatory function

Genome feature analysis reveals that VMRs were preferentially enriched in gene regulatory elements, especially at 2,000 bp upstream regions of genes and the first introns (Fig. 3B and Supplementary Table S5 for counts, frequencies and associated statistics). This pattern is particularly notable for the liver (intron 1 frequency: 23.7 %; promoter frequency: 9.7 %) compared to the whole-genome distribution of CpGs (intron 1 frequency: 14.9 %; promoter frequency: 4.7 %; Supplementary Table S5).

Given their distribution in gene regulatory sequences, and in order to better understand the functional importance of VMRs, we performed a Gene Ontology (GO) enrichment analysis of genes associated with VMRs. Our analysis revealed 20 biologically relevant clusters that can be rooted in 5 larger functional categories related to the neural function, various cellular processes such as transport, signaling and architecture, as well as development and differentiation (Fig. 4, Supplementary Fig. 2 and Supplementary Table S6). In the liver, significant GO terms (p < 0.01) were grouped into clusters related to GABAergic synaptic transmission, negative regulation of signaling pathways, and angiogenesis-linked morphogenesis. These clusters involved between 6 and 88 unique liver-specific genes. In the hypothalamus, the enrichment was more extensive, with 16 GO terms associated with a single cluster and up to 1,649 unique genes in that category alone. Prominent hypothalamic clusters included processes such as MAPK/JNK signaling, regulation of synaptic transmission, and developmental pathways, all of which are relevant to neuronal plasticity and signal integration (Supplementary Table S6). Some functional categories, such as MAPK cascade signaling, were exclusive to the hypothalamus, indicating potential tissue-specific function of VMR-associated genes. Interestingly, many clusters (15/20) were significantly enriched for both tissues, and both tissues were represented in the 5 large functional categories, despite the lower number of GO terms for liver. This result is consistent with the fact that a subset of VMRs is colocalized between the two tissues.

**Fig. 4:**
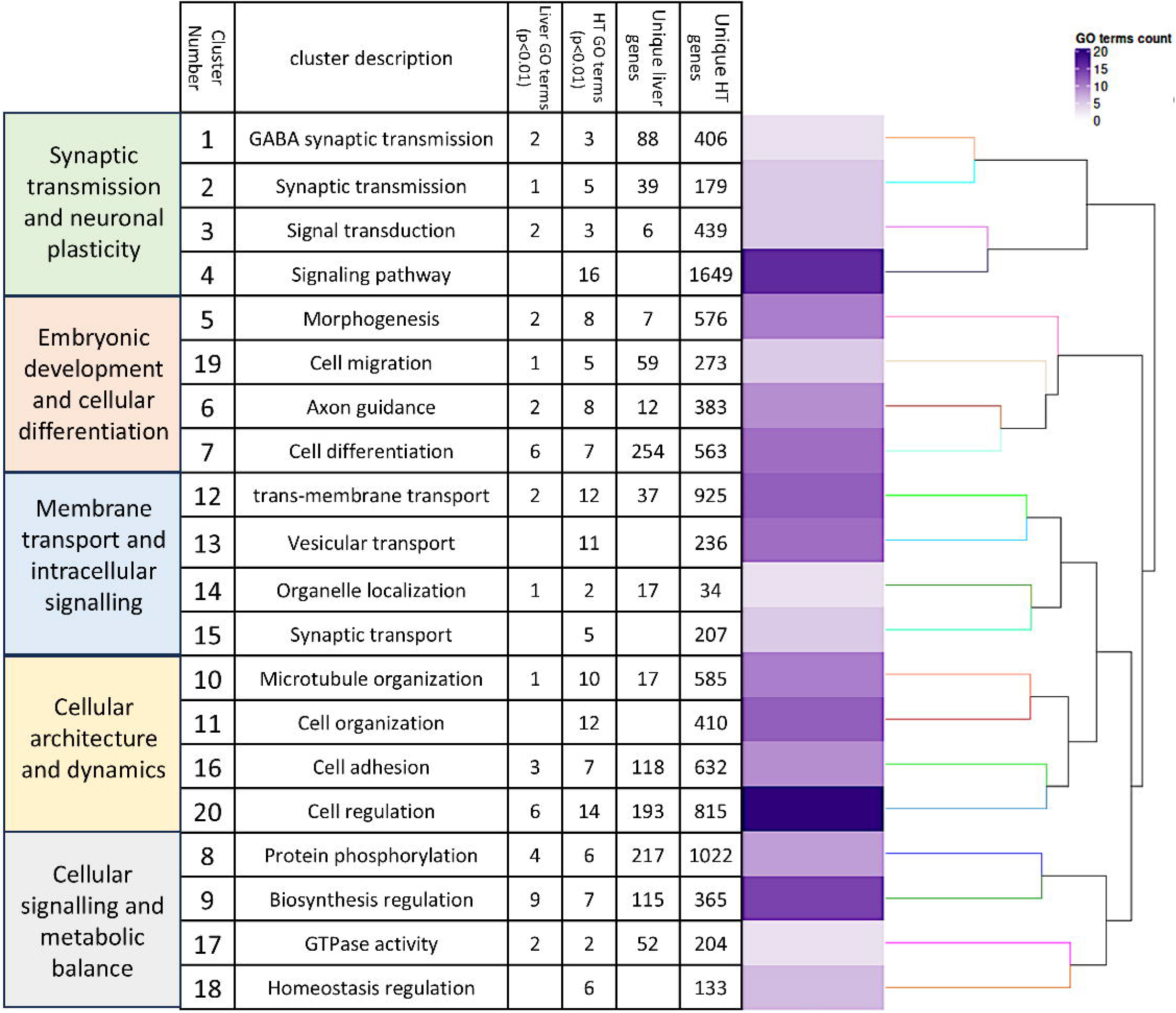
Summary of the Gene Ontology functional analysis of the VMR-associated genes. The clustering heat map plots of the functional sets of gene ontology (GO) terms were obtained using ViSEAGO based on the genes overlapping (at least 1bp) the VMRs. VMRs (Q90) from both tissues were combined in this analysis (for a more detailed view see Supplementary Figure S2). From left to right are shown the five function groups, the 20 functional clusters (cl) with the corresponding heat map (in purple tone) showing GO term counts and a dendrogram on enriched GO terms based on BMA semantic similarity distance and Ward’s clustering criterion. The GO terms defining the clusters and associated statistics are shown in Supplementary Table S6.

To further explore the function of VMRs, we compared the distribution of the VMRs with that of super enhancers (SEs), histone H3 marks related to SEs and ATAC-seq peaks recently published by Salem et al. (Fig. 5 and Supplementary Fig. 3). Strikingly, we found that most brain SEs overlapped with at least one hypothalamic VMR (99.5 %, Jaccard index ∼4,36 %; Fig. 5A), whereas ∼8 % of hepatic SEs overlapped with hepatic VMRs (Jaccard index ∼3,73 %; Fig. 5B). The overlap seemed predominant for tissue-specific SEs, as only 2 % of SEs found in both brain and hepatic tissues overlapped with VMRs found in both hypothalamic and hepatic tissues (Jaccard index ∼1,12 %; Fig. 5C). All overlaps were significant according to a hypergeometric enrichment test using a genomic background of approximately 15.3 million 150 bp windows (based on a 2.3 Gb genome), suggesting a non-random co-localization of VMRs with SEs.

**Fig. 5:**
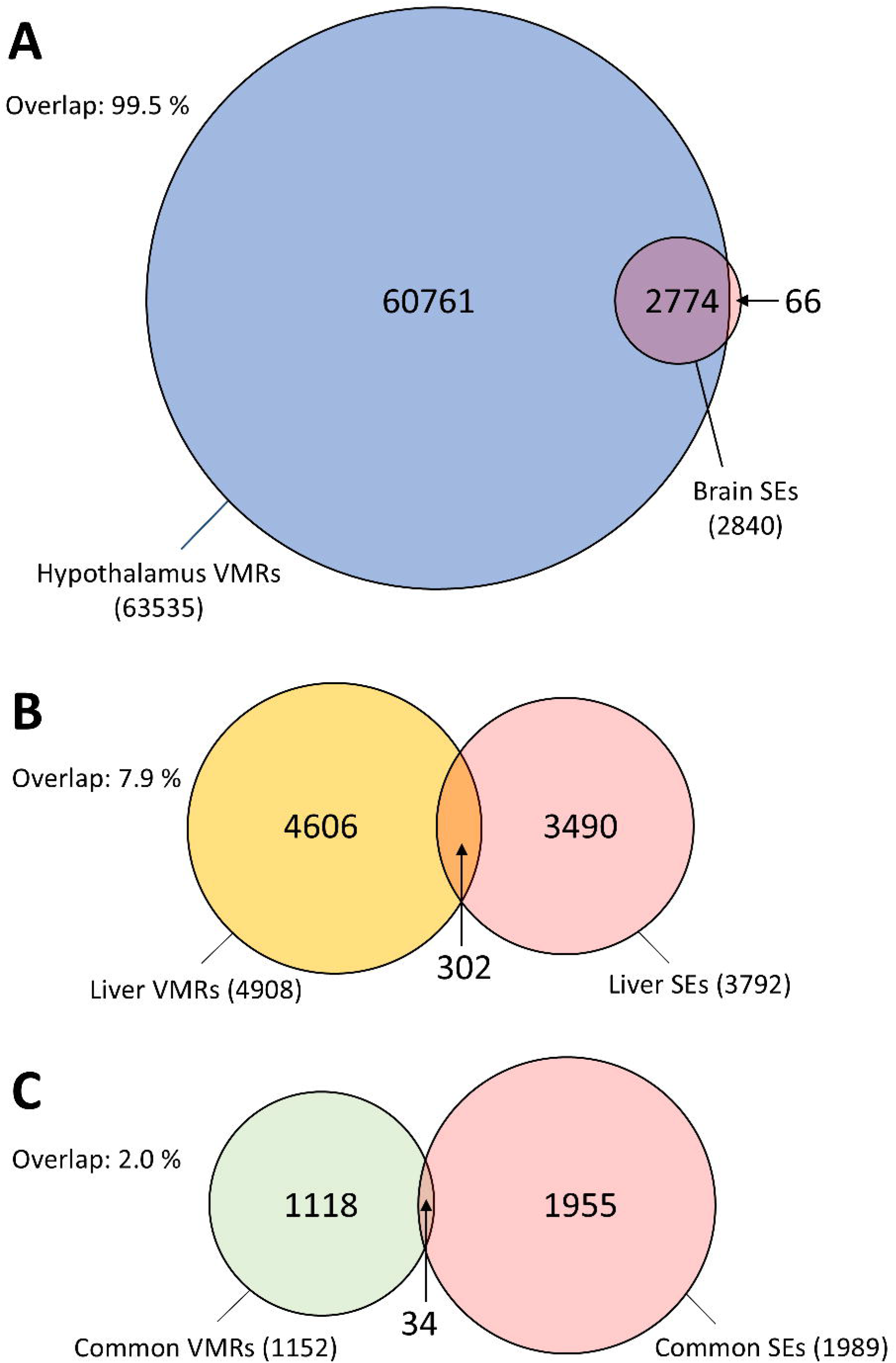
Overlap analysis of VMRs and super-enhancers (SEs) from Salem et al. (2024) in brain and liver tissues. Venn diagrams show that (A) 99.5 % of brain SEs overlap with hypothalamic VMRs, compared with (B) 7.9 % of liver SEs that overlap with liver VMRs and (C) 2.0 % of common SEs that overlap with common VMRs. The numbers of VMRs (Q90) and SEs in each category are indicated.

### Methylation levels of common VMRs show characteristics similar to those of CorSIVs

In mammals, CorSIVs (correlated regions of systemic interindividual variation) have been shown to be VMRs whose DNA methylation levels are consistently correlated across multiple tissues within the same individual [5,21,22]. We sought to determine whether the common VMRs (i.e., not specific to one tissue) identified in trout showed correlated methylation levels within each individual, such as CorSIVs. Indeed, the methylation profiles of common VMRs were correlated in both tissues for a given individual, although this level varied from one VMR to another (Fig. 6A). This was confirmed by a stronger correlation of methylation levels for common VMRs compared with tissue-specific VMRs, especially for the VMRs with the most confidence, i.e. identified at the 99th percentile (Fig. 6B).

**Fig. 6:**
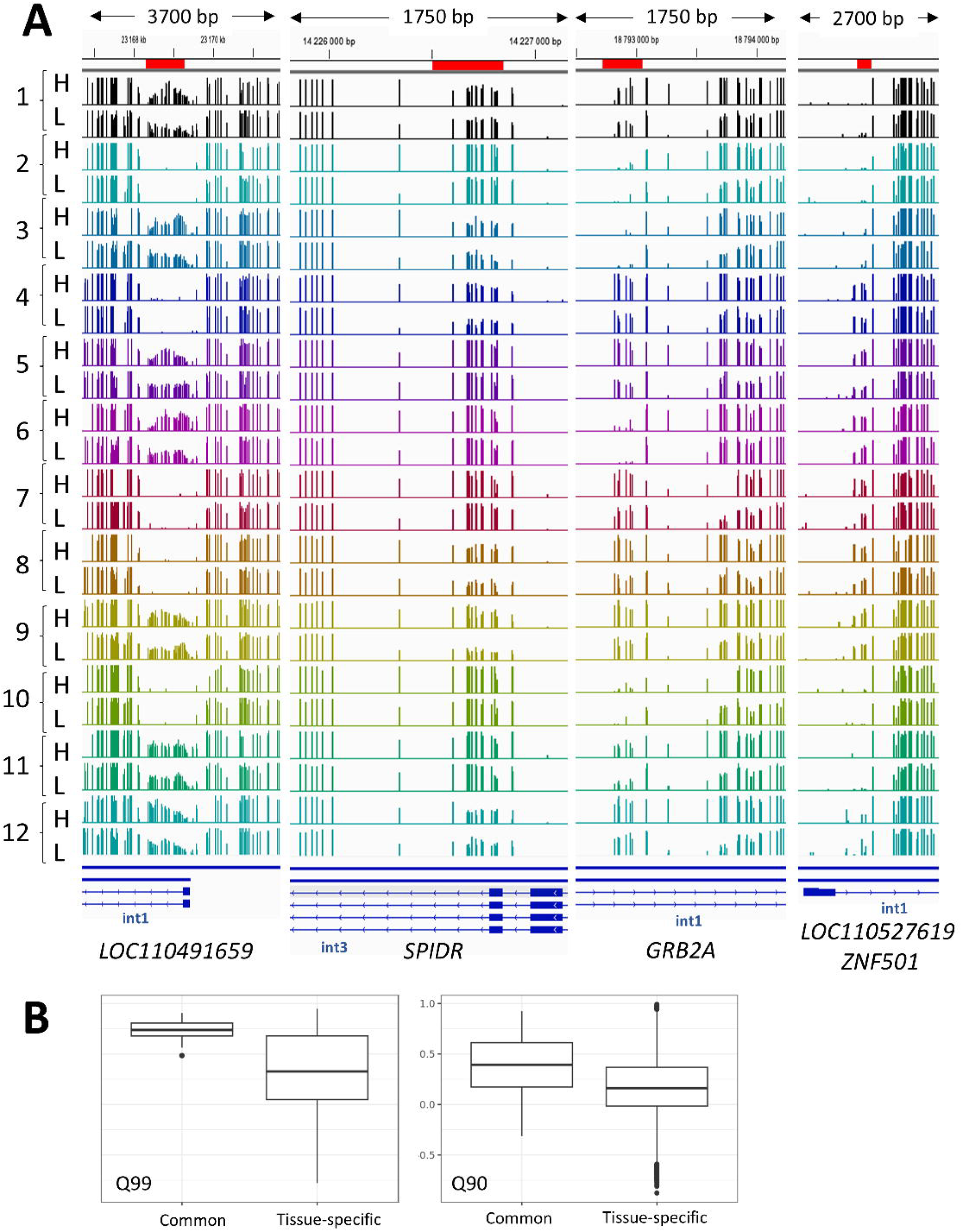
VMRs common to both tissues show correlated levels of methylation. (A) Methylome tracks showing the methylation levels at 4 genomic loci for the 12 individuals. Tracks from the hypothalamus (H) and the liver (L) are grouped by individual (one color per individual). The number of the individual is written on the left. VMR coordinates (are shown in red boxes. The diagram below the tracks shows the relative positions of exons (plain boxes), introns (narrow arrowed lines), and neighboring genes with directional arrows indicating gene orientation. (B) Boxplots showing the correlation of methylation levels for common VMRs and uncommon VMRs (defined by a VMR in one tissue that is not a VMR in the other) at Q90 and Q99.

## Discussion

The study aimed to identify interindividual DNA methylation variation in rainbow trout by analyzing the hepatic and hypothalamic methylomes of isogenic fishes raised in a standard control environment. The selection of parameters was a crucial step in identifying VMRs with confidence. Setting a minimum coverage threshold was necessary to ensure the reliability of methylation calls. A threshold of 10 is often used to balance the need for accurate methylation estimates while maintaining a sufficient number of regions for analysis. This threshold helps to avoid biases due to low coverage, which can lead to inaccurate methylation levels [23]. Requiring the minimum coverage (≥ 10) to be present in at least 50 % of the samples ensured that the regions analyzed are consistently covered across the dataset. This helped in reducing variability and increasing the confidence in the methylation calls while maintaining a balance between the number of regions and the quality of the data. Moreover, MAD was preferred over SD (standard deviation) in scenarios with a small number of samples because it is less sensitive to outliers. This makes it a more robust measure of variability, ensuring that the analysis is not unduly influenced by extreme values [24]. While our study uses the concept of variability (MAD) to define VMRs, some human and mouse studies employed information-theoretic methods like methylation stochasticity or entropy to capture other aspects of variability not detectable by mean-based analyses [25,26]. Methylation entropy changes were shown to contribute comparably to or more than mean methylation changes in shaping the dynamic epigenetic landscape during mouse development [25]. Therefore, focusing solely on variance might only capture one dimension of the complex methylation variability landscape.

The choice of parameters such as the maximum distance between VMCs and the minimum number of VMCs to define VMRs was based on the need to identify biologically relevant regions. These parameters are often determined empirically and can vary depending on the specific goals of the study. Altogether, the selection of parameters that minimized the distance between VMCs while ensuring a sufficient number of VMRs (100 bp and 5 VMCs per VMR) resulted in the identification of 139 / 4,908 (Q99 / Q90) and 814 / 63,535 VMRs for the liver and hypothalamus, respectively. This approach allowed for a detailed examination of methylation variation within these tissues. The confirmation of a VMR located in the first intron of the LOC110491659 gene through bisulfite conversion and Sanger sequencing validated the method’s accuracy. The absence of *cis* polymorphisms further supported the epiallelic nature of the identified VMR. Overall, this analysis confirms the existence of VMRs in rainbow trout. Interestingly, more VMRs were identified in the liver than in the hypothalamus (about 6 times more at Q99 and 12 times more at Q90). One possible explanation is that 27.3 % more cytosines were analyzed in the hypothalamus than in the liver (Supplementary Tables S1 and S2). This difference could also be explained by the higher cellular heterogeneity of the hypothalamic (due to sampling variation or the intrinsic nature of the tissue) compared to the liver [27,28], despite a previous report suggesting that methylation variation at hyper-variable CpGs was not driven by cell heterogeneity effects [29]. However, another study in cattle showed that the number of highly variable CpGs was similarly higher in the anterior pituitary, a gland just below the hypothalamus, than in leukocytes under control conditions [5]. This observation was attributed primarily to cell type heterogeneity within the tissue samples. To clarify this point, future studies should focus on a single cell-type to avoid the potential confounding effect of different methylation profiles resulting from different cell types.

A key aspect of our study design on rainbow trout was the use of an isogenic fish line, selected to minimize genetic variation and thus likely isolate intrinsic epigenetic variability [30]. This contrasts with the approach and findings in human and cattle studies, which consistently highlight the strong influence of genetic variation on interindividual DNA methylation patterns at VMRs [4,21,22,29,31]. Human studies show that genetic influences on VMR methylation can be much stronger than at other CpG sites interrogated by standard arrays [21]. Our finding that a validated VMR in an intron of LOC110491659 was not associated with *Cis* polymorphisms is expected because of the genetic uniformity within an isogenic line, which effectively removed a major source of VMR formation observed in genetically diverse human and cattle populations.

Analysis of the distribution of VMRs in relation to genomic features suggests a function in the regulation of gene expression, in particular around the TSS at the 1^st^ intron, and in the promoter for liver. Strikingly, a significant overlap between VMRs and markers of open chromatin (H3K4me1 and ATACseq peaks) defining super-enhancers as identified in [15] was found notably in the hypothalamus, reinforcing the hypothesis of a contribution to gene regulation. In humans and mice, VMRs were also shown to be often located near gene regulatory elements, such as promoters and enhancers, colocalizing with enhancer histone modifications [29,31,32]. However, one study reported that the majority (88 %) of human VMRs were located more than 5 kb away from annotated TSSs [33], suggesting they might function as distal regulatory elements. In our analysis, no clear enrichment of VMRs in intergenic regions (“other” in Fig 3B) was observed. Furthermore, in cattle, a subset of VMRs was enriched at TSSs, but not at intronic or intergenic regions [22]. Altogether, despite a few discrepancies that may be caused by the different statistical methods or experimental designs, the preferential distribution of VMRs near gene regulatory elements in rainbow trout suggests a conserved mechanism of epigenetic regulation across vertebrates.

Our observation that VMRs that colocalize in both tissues display a strong intraindividual correlation in methylation levels, particularly for high-confidence regions (Fig. 6A–B), provides important insight into the developmental origins and stability of epigenetic variation. This finding is particularly striking given the likely absence of underlying genetic polymorphisms in the isogenic line used in this study, indicating that the methylation differences are likely to arise from non-genetic factors established early during development. This pattern is reminiscent of correlated regions of systemic interindividual variation (CoRSIVs) previously described in humans as genomic regions where DNA methylation varies between individuals but remains highly consistent across tissues within the same individual [21,29]. CoRSIVs have been shown in humans and cattle [5,21,22] and are believed to originate during the peri-implantation phase, following the wave of global epigenetic reprogramming in the early embryo, and are thought to represent stable epigenetic “set points” established before the onset of lineage specification. Our findings extend this concept to a fish model and suggest that shared VMRs may serve a similar role, acting as systemically stable epigenetic marks. The higher methylation concordance observed in shared versus tissue-specific VMRs, especially in the most confidently identified ones, supports the notion that these regions may be established during early embryogenesis, prior to the divergence of tissue-specific methylation landscapes. The presence of such regions may reflect early stochastic events, maternal environment effects, or epigenetic noise, which become fixed and propagated through subsequent cell divisions across lineages. Such regions may contribute to systemic variation in gene regulation, influencing pathways such as immune function, metabolism, and neural development. In contrast, tissue-specific VMRs, which show lower concordance between tissues, likely reflect post-differentiation influences, including tissue-specific transcription factor activity and environmental exposures. The distinction between shared and tissue-specific VMRs underscores the layered nature of the epigenome, with some components reflecting early, systemic programming and others representing dynamic, tissue-dependent regulation. Taken together, although this study focuses on just two tissues, our results support a model in which a subset of common VMRs, analogous to CorSIVs, are epigenetically determined during early development and contribute to stable interindividual differences across tissues. These regions may offer valuable insights into epigenetic memory and phenotypic variation, especially in the context of disease susceptibility and developmental biology. The absence of sequence polymorphisms in the isogenic line of rainbow trout reinforces the idea that these epigenetic variations are intrinsically determined and may play a role in developmental programming and phenotypic plasticity.

The functional analysis revealed 20 GO terms clusters that can be further grouped into 5 larger categories. Many of these functions related to core biological processes are highly conserved across vertebrates, including fish and mammals. These functions might indicate a contribution of VMRs to processes related to growth, immune response, environmental adaptation, and neural plasticity. Interestingly, a majority of clusters contain enriched GO terms from both liver and brain. Therefore, VMRs are linked to a broad range of biological functions in a largely non–tissue-specific manner. In general, GO terms were related to the neurotransmission and synaptic signaling (e.g., GABAergic, glutamatergic, chemical synaptic transmission) and the neural regulation, suggesting a strong enrichment of neurobiological functions, particularly in the hypothalamus, and possibly overlapping regulatory functions in the liver. Strikingly, the presence of more than 1,600 unique genes in a single hypothalamic cluster (number 4, related to synaptic transmission and neuronal plasticity) suggests that DNA methylation may contribute to a high degree of regulatory plasticity in the brain. These findings reinforce the idea that VMRs are not randomly distributed but are instead positioned close to genes with key biological pathways, potentially linking environmental inputs to long-term functional outcomes in a tissue-dependent manner. VMRs in humans and cattle also cover genes involved in brain function and metabolic processes [5,26,33], suggesting a conserved role of VMRs in regulating core biological processes, including neural but also metabolic functions, across vertebrates. In particular, cluster 9 is related to hepatic lipid metabolism. The GO terms related to diacylglycerol and acyl-CoA metabolic processes were found enriched in the liver and suggest that VMR may affect genes related to energy storage and mobilization. Similarly, the ubiquinone biosynthetic process plays a role in mitochondrial respiration and oxidative phosphorylation, and is crucial for energy production. Serotonin biosynthetic process, while often associated with neural function, also occurs in peripheral tissues like the liver, where serotonin modulates hepatic regeneration and vascular tone. Thus, VMRs may impact metabolic and regulatory functions essential to hepatic physiology. Notably, we found that CPT1a-like liver isoform (LOC110491659) has a CorSIV-like VMR in the first intron of the gene. CPT1a is a key regulator of mitochondrial β-oxidation, and its expression influences lipid catabolism and energy homeostasis [34]. Variable methylation of this locus, probably under the influence of development or the environment, may therefore lead to interindividual differences in energy use, growth, performance or feed efficiency. Of interest, the methylation status of the CPT1a-like VMR was either methylated or unmethylated compared to other loci that showed greater variation (Fig. 1 and 6), suggesting an ON/OFF mechanism. This is reminiscent of previously described metastable epiallele such as the human *POMC* VMR [32] and the mouse *A*^*vy*^ locus [35]. It remains to be established whether CPT1a-like epigenetic variation has a programming effect on metabolism as shown for the other metastable epialleles, nevertheless these findings suggest that CPT1a-like is a relevant candidate for studying the epigenetic basis of metabolic plasticity in fish.

The identification and characterization of VMRs in rainbow trout isogenic line raised in a single condition provide valuable insights into the regulatory roles of DNA methylation in gene expression and biological processes. The preferential distribution of VMRs near gene regulatory elements, their overlap with super enhancers, and their establishment during early embryogenesis underscore their conservation among vertebrates and their significance in developmental and physiological contexts. Further studies are required to explore the functional consequences of these VMRs on the expression of the associated genes, in different environmental conditions and developmental stages, potentially uncovering new mechanisms of fine-tuning of gene expression in rainbow trout transferable to other species.

## List of abbreviations

ATAC: Assay for Transposase-Accessible Chromatin
A^vy^: Agouti viable yellow
BMA: Best Match Average
Bp: Base pair
CPT1A: Carnitine Palmitoyltransferase 1A
CpG: Cytosine-phosphate-Guanine dinucleotide
DNA: Deoxyribonucleic acid
gDNA: Genomic
DNA GO: Gene Ontology
H3K4me1: Histone H3 lysine 4 mono-methylation
HMW: High molecular weight (DNA)
HT: Hypothalamus
IAP: Intracisternal A-particle (retrotransposon)
IGV: Integrative Genomics Viewer
MAD: Median absolute deviation
PBAT: Post-Bisulfite Adaptor Tagging
PCR: Polymerase Chain Reaction
POMC: Pro-opiomelanocortin
RNA: Ribonucleic acid
R: R Project for Statistical Computing
SE: Super enhancer
TSS: Transcription start site
UTR: Untranslated region
VMR: Variably methylated region
VMC: Variably methylated cytosine
WGBS: Whole-genome bisulfite sequencing

## Declarations

### Ethics approval and consent to participate

The experiments were in strict accordance with EU legal frameworks related to the protection of animals used for scientific research (Directive 2010/63/EU) and according to the National Guidelines for Animal Care of the French Ministry of Research (decree n°2013-118, February 1st, 2013). In agreement with ethical committee “Comité d’Éthique Aquitaine Poissons Oiseaux” (C2EA-73), the experiment reported here does not need approval by a specific ethical committee since it implies only classical rearing practices with all diets used in the experimental formulated to cover all the nutritional requirements of rainbow trout [36].

### Consent for publication

Not applicable

### Availability of data and materials

All data supporting the findings in this study are available within the article and its supplementary information files. The raw WGBS data is available at the European Nucleotide Archive (ENA) under the accession number PRJEB86744. Correspondence and material requests should be addressed to Vincent Coustham.

### Competing interests

The authors declare that they have no competing interests.

### Funding

This study was funded by INRAE IB22 API PHASE “EVE” and by the project Initiative E2S MIRA - Fédération de Recherche sur les Milieux et Ressources Aquatiques.

### Authors’ contributions

VC, FT and GL designed the experiments. DL provided the isogenic line. VC, AP, LB, FT, JR, LM and SP performed the experiments. VC, GL, AB and BP analyzed the results. BP managed and submitted the data to the ENA repository. VC, GL and AB wrote the manuscript. All authors read and approved the final manuscript.

## Acknowledgements

The authors are grateful to the members of the NuMéA unit at INRAE for assistance with sample collection and to the staff of the INRAE fish facilities at Lees-Athas, Donzacq and PEIMA for animal care and management. The authors thank Audrey Laurent (INRAE, LPGP, France) for critical reading of the manuscript. This work also benefited from the IT infrastructure of the ISLANDe platform, and particularly a computing cluster financed by the European Regional Development Fund n°159037.

## Supplementary Figure Legends

**Supplementary Fig. S1:** Coverage analysis relative to significance thresholds. Number of positions remaining when removing those for which the coverage is between 5 and 12 for 10 to 90 % of the samples.

**Supplementary Fig. S2:** Complete Gene Ontology functional analysis of the VMR-associated genes (Q90). The clustering heat map plots of the functional sets of gene ontology (GO) terms were obtained using ViSEAGO. From left to right are shown the enriched GO terms, a heat map (in red tones) showing –log10 p-value of enrichment test for liver, a heat map (in red tones) showing –log10 p-value of enrichment test for hypothalamus, a heat map (in purple tone) showing information content and a dendrogram on enriched GO terms based on BMA semantic similarity distance and Ward’s clustering criterion. F: liver; HT: hypothalamus; IC: Information Content.

**Supplementary Fig. S3:** Count analysis of the number of histone H3 marks (H3K4me1, H3K4me3, H3K27Ac and H3K27me3), ATAC-seq peaks and super enhancer regions (SE) that overlap with VMRs (Q90) for (A) hypothalamus and (B) liver. ChIP-seq, ATAC-seq and SE data was taken from Salem et al., 2024. The percentage of overlapping marks, ATAC-seq peaks and SEs with VMRs are shown below.

## Supplementary Table Legends

**Supplementary Table S1:** Summary of the MultiQC report on the sequencing data from liver tissue following quality/adapter trimming and Bismark analyses.

**Supplementary Table S2:** Summary of the MultiQC report on the sequencing data from hypothalamus tissue following quality/adapter trimming and Bismark analyses.

**Supplementary Table S3:** Number of CpGs analyzed and % relative to the total number of genomic CpGs based on the coverage threshold (5 to 12) and the percentage of samples (10 to 90 %).

**Supplementary Table S4:** VMR parameters (columns A-C) and associated results for liver (columns D-G) and hypothalamus (columns H-K)

**Supplementary Table S5:** Genome feature analysis results.. The count number represents the number of CpGs covered by VMRs for each feature, and the frequency represents the proportion of each feature’s count relative to all features in the genome. “Promoter” is defined as the region 2,000 bp upstream of the Transcription Start Site, and “downstream” is defined as the region 1,000 bp downstream from the Transcription End Site. P-values correspond to a Fisher’s exact test (one-sided, greater alternative) based on the VMRs CpG content. “Genome” refers to the number and frequency of genomic features of all CpGs in the genome.

**Supplementary Table S6:** Gene Ontology (GO) enrichment results associated with VMRs (Q90) in liver and hypothalamus (HT), obtained using the ViSEAGO R package [17]. Each line corresponds to a single GO term. Information content (IC) is computed as the negative log probability of occurrence of the term in a set of GO terms. NA: Not Available.

**Supplementary Table S7:** List of the primers used in the study.

## Notes

https://www.ebi.ac.uk/ena/browser/view/PRJEB86744

### Competing Interest Statement

The authors have declared no competing interest.

