## Supplementary Figures for "Characterization of interindividual DNA methylation variability in rainbow trout (*Oncorhynchus mykiss*)"

### Supplementary Figure S1

**A** – Number of positions

**B** – Number of regions

**C** – Median region size

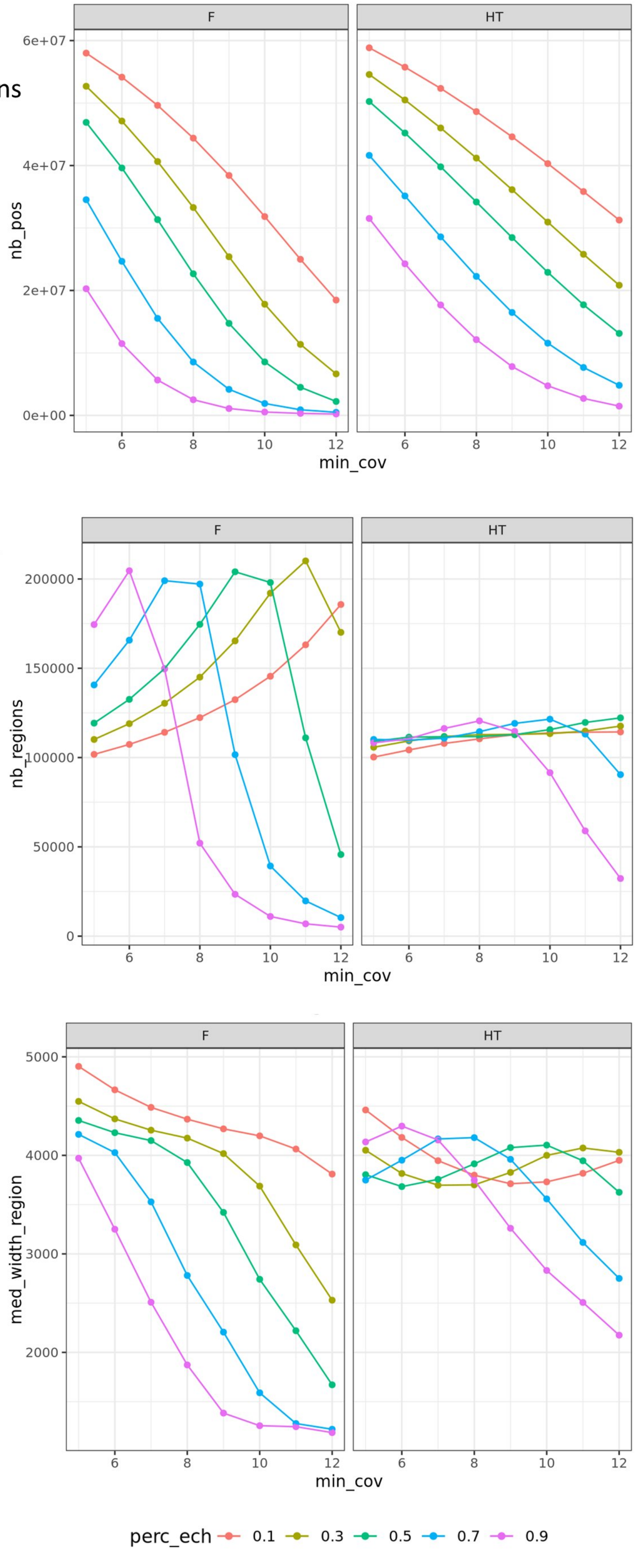

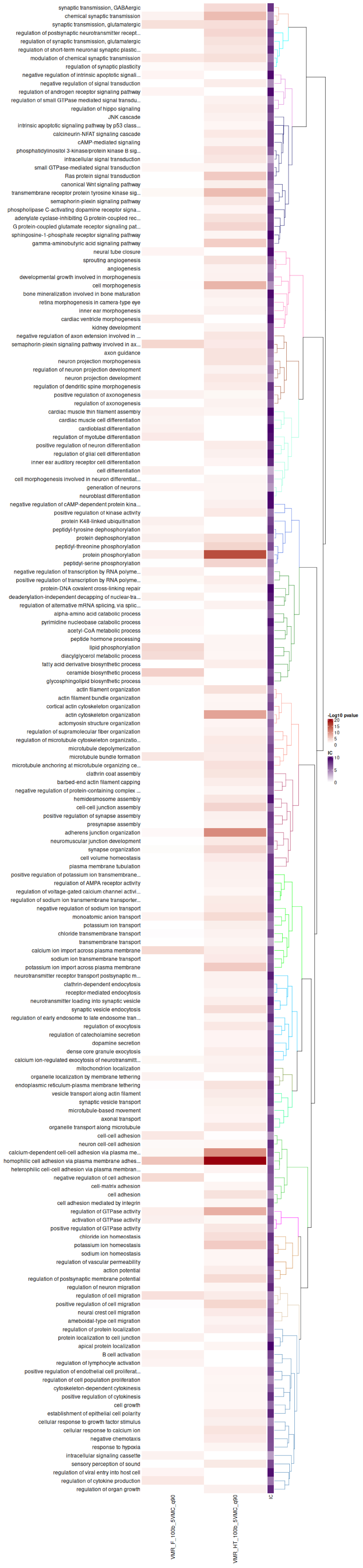
synaptic transmission, GABAergic
chemical synaptic transmission
synaptic transmission, glutamatergic
regulation of postsynaptic neurotransmitter recept...
regulation of synaptic transmission, glutamatergic
regulation of short-term neuronal synaptic plastic...
modulation of chemical synaptic transmission
regulation of synaptic plasticity
negative regulation of intrinsic apoptotic signali...
negative regulation of signal transduction
regulation of androgen receptor signaling pathway
regulation of small GTPase mediated signal transdu...
regulation of hippo signaling
JNK cascade
intrinsic apoptotic signaling pathway by p53 class...
calcineurin-NFAT signaling cascade
cAMP-mediated signaling
phosphatidylinositol 3-kinase/protein kinase B sig...
intracellular signal transduction
small GTPase-mediated signal transduction
Ras protein signal transduction
canonical Wnt signaling pathway
transmembrane receptor protein tyrosine kinase sig...
semaphorin-plexin signaling pathway
phospholipase C-activating dopamine receptor signa...
adenylate cyclase-inhibiting G protein-coupled rec...
G protein-coupled glutamate receptor signaling pat...
sphingosine-1-phosphate receptor signaling pathway
gamma-aminobutyric acid signaling pathway
neural tube closure
sprouting angiogenesis
angiogenesis
developmental growth involved in morphogenesis
cell morphogenesis
bone mineralization involved in bone maturation
retina morphogenesis in camera-type eye
inner ear morphogenesis
cardiac ventricle morphogenesis
kidney development
negative regulation of axon extension involved in ...
semaphorin-plexin signaling pathway involved in ax...
axon guidance
neuron projection morphogenesis
regulation of neuron projection development
neuron projection development
regulation of dendritic spine morphogenesis
positive regulation of axonogenesis
regulation of axonogenesis
cardiac muscle thin filament assembly
cardiac muscle cell differentiation
cardioblast differentiation
regulation of myotube differentiation
positive regulation of neuron differentiation
regulation of glial cell differentiation
inner ear auditory receptor cell differentiation
cell differentiation
cell morphogenesis involved in neuron differentiat...
generation of neurons
neuroblast differentiation
negative regulation of cAMP-dependent protein kina...
positive regulation of kinase activity
protein K48-linked ubiquitination
peptidyl-tyrosine dephosphorylation
protein dephosphorylation
peptidyl-threonine phosphorylation
protein phosphorylation
peptidyl-serine phosphorylation
negative regulation of transcription by RNA polyme...
positive regulation of transcription by RNA polyme...
protein-DNA covalent cross-linking repair
deadenylation-independent decapping of nuclear-tra...
regulation of alternative mRNA splicing, via splic...
alpha-amino acid catabolic process
pyrimidine nucleobase catabolic process
acetyl-CoA metabolic process
peptide hormone processing
lipid phosphorylation
diacylglycerol metabolic process
fatty acid derivative biosynthetic process
ceramide biosynthetic process
glycosphingolipid biosynthetic process
actin filament organization
actin filament bundle organization
cortical actin cytoskeleton organization
actin cytoskeleton organization
actomyosin structure organization
regulation of supramolecular fiber organization
regulation of microtubule cytoskeleton organizatio...
microtubule depolymerization
microtubule bundle formation
microtubule anchoring at microtubule organizing ce...
clathrin coat assembly
barbed-end actin filament capping
negative regulation of protein-containing complex ...
hemidesmosome assembly
cell-cell junction assembly
positive regulation of synapse assembly
presynapse assembly
adherens junction organization
neuromuscular junction development
synapse organization
cell volume homeostasis
plasma membrane tubulation
positive regulation of potassium ion transmembrane...
regulation of AMPA receptor activity
regulation of voltage-gated calcium channel activi...
regulation of sodium ion transmembrane transporter...
negative regulation of sodium ion transport
monoatomic anion transport
potassium ion transport
chloride transmembrane transport
transmembrane transport
calcium ion import across plasma membrane
sodium ion transmembrane transport
potassium ion import across plasma membrane
neurotransmitter receptor transport postsynaptic m...
clathrin-dependent endocytosis
receptor-mediated endocytosis
neurotransmitter loading into synaptic vesicle
synaptic vesicle endocytosis
regulation of early endosome to late endosome tran...
regulation of exocytosis
regulation of catecholamine secretion
dopamine secretion
dense core granule exocytosis
calcium ion-regulated exocytosis of neurotransmitt...
mitochondrion localization
organelle localization by membrane tethering
endoplasmic reticulum-plasma membrane tethering
vesicle transport along actin filament
synaptic vesicle transport
microtubule-based movement
axonal transport
organelle transport along microtubule
cell-cell adhesion
neuron cell-cell adhesion
calcium-dependent cell-cell adhesion via plasma me...
homophilic cell adhesion via plasma membrane adhes...
heterophilic cell-cell adhesion via plasma membran...
negative regulation of cell adhesion
cell-matrix adhesion
cell adhesion
cell adhesion mediated by integrin
regulation of GTPase activity
activation of GTPase activity
positive regulation of GTPase activity
chloride ion homeostasis
potassium ion homeostasis
sodium ion homeostasis
regulation of vascular permeability
action potential
regulation of postsynaptic membrane potential
regulation of neuron migration
regulation of cell migration
positive regulation of cell migration
neural crest cell migration
ameboidal-type cell migration
regulation of protein localization
protein localization to cell junction
apical protein localization
B cell activation
regulation of lymphocyte activation
positive regulation of endothelial cell proliferat...
regulation of cell population proliferation
cytoskeleton-dependent cytokinesis
positive regulation of cytokinesis
cell growth
establishment of epithelial cell polarity
cellular response to growth factor stimulus
cellular response to calcium ion
negative chemotaxis
response to hypoxia
intracellular signaling cassette
sensory perception of sound
regulation of viral entry into host cell
regulation of cytokine production
regulation of organ growth
VMR_F_100b_5VMC_q90
VMR_HT_100b_5VMC_q90
IC
-Log10 pvalue
20
15
10
5
0
IC
10
5
0

**Supplementary Figure S3**

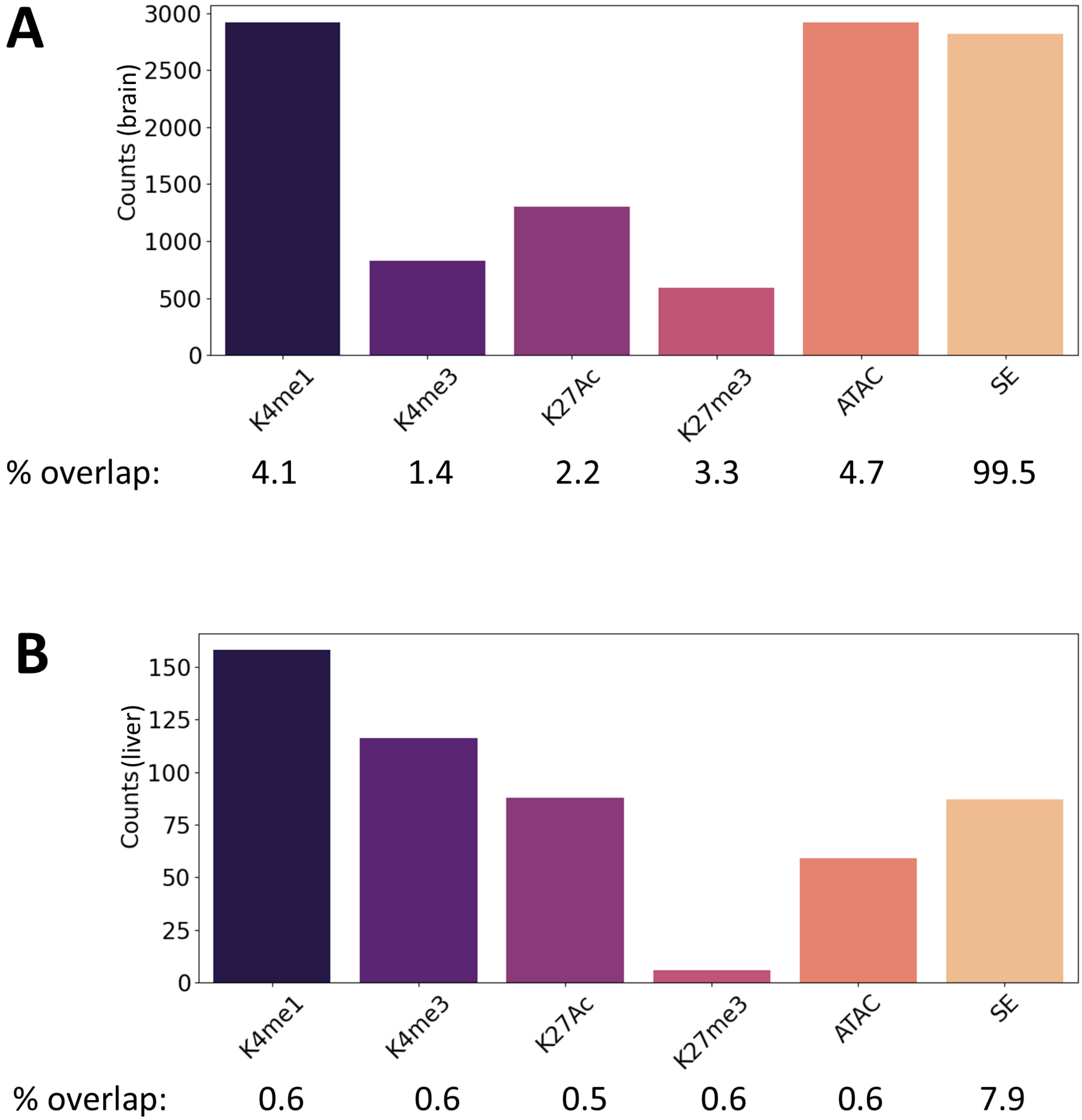
